# From neurons to voxels - repetition suppression is best modelled by local neural scaling

**DOI:** 10.1101/170498

**Authors:** Arjen Alink, Hunar Abdulrahman, Richard N. Henson

## Abstract

Inferring neural mechanisms from functional magnetic resonance imaging (fMRI) is challenging because the fMRI signal integrates over millions of neurons. One approach is to compare computational models that map neural activity to fMRI responses, to see which best predicts fMRI data. We used this approach to compare four possible neural mechanisms of fMRI adaptation to repeated stimuli (scaling, sharpening, repulsive shifting and attractive shifting), acting across three domains (global, local and remote). Six features of fMRI repetition effects were identified, both univariate and multivariate, from two independent fMRI experiments. After searching over parameter values, only the local scaling model could simultaneously fit all data features from both experiments. Thus fMRI stimulus repetition effects are best captured by down-scaling neuronal tuning curves in proportion to the difference between the stimulus and neuronal preference. These results emphasize the importance of formal modelling for bridging neuronal and fMRI levels of investigation.

---

Inferring the properties of neurons from gross measurements like functional magnetic resonance imaging (fMRI) is notoriously difficult. Because the fMRI signal integrates over millions of neurons, such inferences represent an inverse problem with no unique solution. However, progress can be made by comparing different “forward” models that formalize the mapping from neural activity to fMRI responses. This problem applies for example when trying to understand the phenomenon of reduced fMRI responses associated with repeated stimulation – referred to as “fMR adaptation” or “repetition suppression” (Ashida et al., 2007; Cavina-Pratesi et al., 2010; Chong et al., 2008; Fang et al., 2005; Fox et al., 2009; Grill-Spector et al., 2006; Henson, 2016; Kar and Krekelberg, 2016; Krekelberg et al., 2006; Winston et al., 2004) (Winston et al., 2004). These fMRI reductions have been attributed to a range of different neural mechanisms, such as “sharpening”, whereby repetition is thought to narrow neural tuning-curves (Gotts et al., 2012; Henson and Rugg, 2003; Weiner et al., 2010; Wiggs and Martin, 1998). Here we address this problem by implementing a range of “forward models” that map from neuronal firing to fMRI signals, and comparing their predictions to key empirical features of fMRI repetition effects.

A previous study by Weiner and colleagues (Weiner et al., 2010) formally modelled the relationship between neuron-level and voxel-level repetition effects. This study was an important demonstration that neural scaling and neural sharpening can reproduce similar effects of repetition on the mean fMRI response across voxels (a “univariate” response), but did not consider the effect of repetition on patterns of fMRI responses across voxels (a “multivariate” response). Indeed, a recent study of multi-voxel patterns by Kok et al. (Kok et al., 2012) showed that prediction of upcoming stimuli (which may also arise from repetition) improved classification of those stimuli, and this was used to support the idea of neural sharpening. However, neither study considered how the effects of repetition on a neural population depend on the difference between a stimulus property and the preference of that neural population for that property. In addition, Kok et al. (Kok et al., 2012) did not specifically test whether improved stimulus classification could also be explained by neural mechanisms other than sharpening. Here we overcome these limitations by considering a wider range of neural models and by evaluating a larger combination of both univariate and multivariate fMRI data features. This enables us to determine, for example, whether repetition effects on fMRI pattern information are specific to neural sharpening.

Our forward models are based on four mechanisms associated with neuronal adaptation, each of which is grounded in previous neurophysiological studies: 1) scaling, where adaptation reduces response amplitude (Ringach and Bredfeldt, 2002; Swindale, 1998), 2) sharpening, where adaptation tightens tuning-curves (Kar and Krekelberg, 2016), 3) repulsive shifting, where the peak of tuning-curves moves away from the adapting stimulus (Dragoi et al., 2001) and 4) attractive shifting, where the peak moves towards the adapting stimulus (Bachatene et al., 2015; Jeyabalaratnam et al., 2013). Each of these basic mechanisms is applied across three domains: 1) global, where all tuning-curves in a voxel are affected, 2) local, where tuning-curves close to the adapting stimulus are affected most, and 3) remote, where tuning-curves close to the adapting stimulus are affected least. Figure 1 illustrates the twelve models resulting from crossing these four mechanisms and three domains. Tuning-curves are modelled by either Gaussian or von Mises distributions, and each model has only 2-3 free parameters: 1) the width of tuning-curves (*σ*), 2) the amount of adaptation (*a*) and, for non-global models, the extent of the domain of adaptation (*b*).

**Figure 1.**
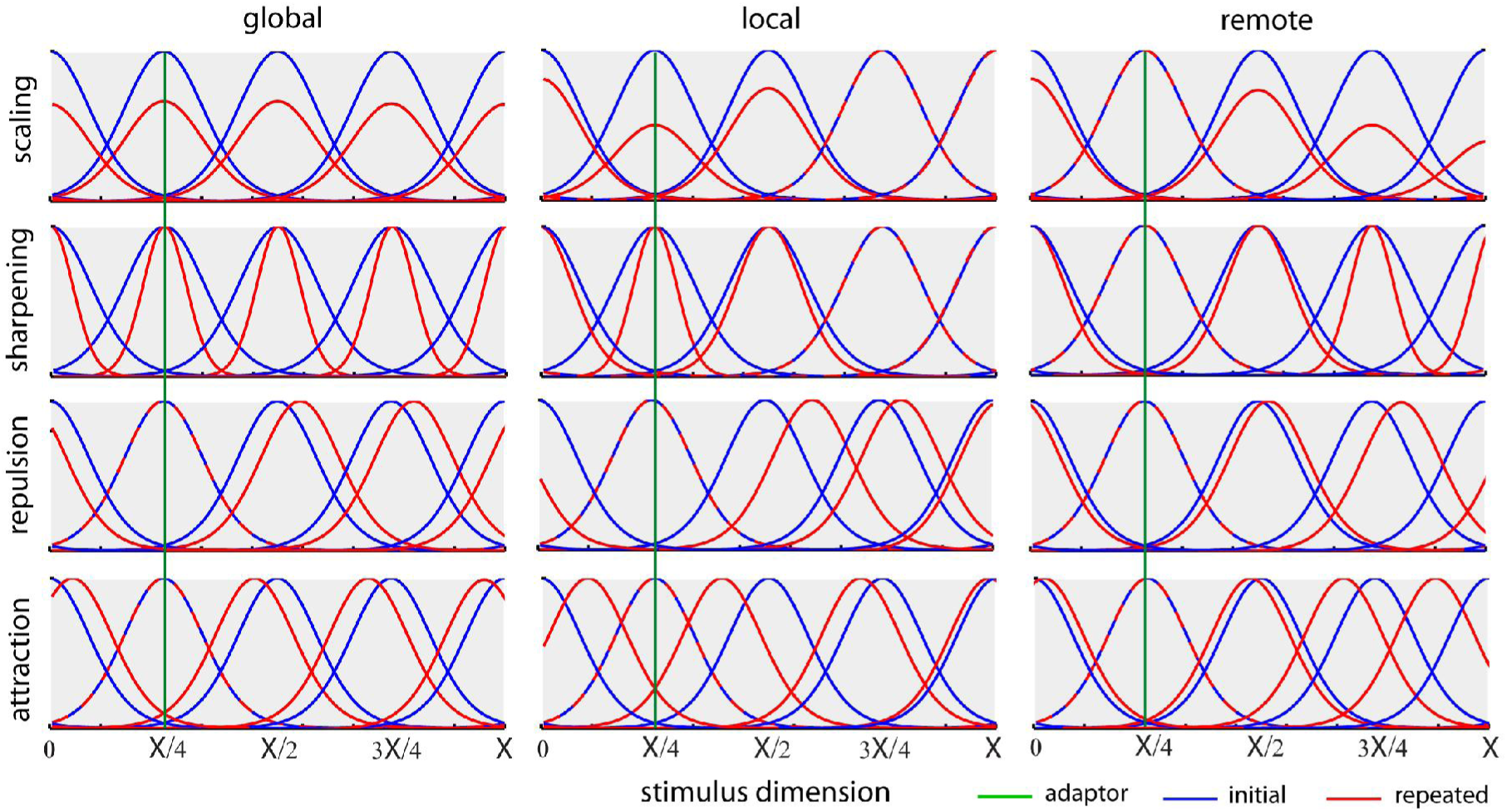
Example tuning curves along a stimulus dimension (ranging from 0 to X), both before (blue) and after (red) repetition of a single stimulus (with value X/4, shown by green line) according to the twelve different neural models of repetition suppression, created by crossing four mechanisms (rows) with three domains (columns). For illustrative purposes, only five neural populations are shown, equally-spaced along the stimulus dimension. Note that this figure illustrates the Gaussian tuning-curves used for Experiment 1 (see Supplemental Figure 1 for illustration of von Mises tuning-curves used for Experiment 2).

We compared the fMRI repetition effects predicted by these models to real fMRI data from two different experiments. Each experiment provided data for two stimulus-classes across multiple voxels within a single region-of-interest (ROI) in the brain. In Experiment 1, the data were responses from voxels in the right fusiform face area (FFA) to initial and repeated brief presentations of face and scrambled face stimuli; in Experiment 2, the data were responses in early visual cortex (V1) to initial and repeated sustained presentations of gratings with one of two orthogonal orientations. For each experiment, we examined how repetition affected six data features: 1) Mean Amplitude Modulation (**MAM**), the traditional univariate measure of repetition suppression, averaged across all voxels in the ROI; 2) Within-class Correlation (**WC**), the mean correlation of multivariate patterns across voxels between all pairs of stimuli of the same class; 3) Between-class Correlation (**BC**), the mean pattern correlation between all pairs of stimuli from different classes; 4) Classification Performance (**CP**), here operationalized as the difference between WC and BC, but validated by a Support Vector Machine (see Methods); 5) Amplitude Modulation by Selectivity (**AMS**), where the repetition-related change in amplitude is binned according to the selectivity of each voxel (defined as the T-value when contrasting the two stimulus classes); and 6) Amplitude Modulation by Amplitude (**AMA**), where the repetition-related amplitude change was binned according to the overall amplitude of each voxel (averaged across initial and repeated presentations). Some of these data features differed across our two experiments, presumably owing to important differences in ROI, stimulus type, stimulus duration, etc. To foreshadow our results, while no data feature on its own was diagnostic of a specific neural model, only the local scaling model could simultaneously reproduce all six data features using the same parameter values, and this was the case for both experiments, despite their different paradigms.

## Results

### Empirical results

The two paradigms are illustrated in Figure 2. Experiment 1 measured the impulse response to brief presentation (<1s) of unfamiliar faces and scrambled faces that repeated immediately on 50% of occasions, in a randomly-intermixed, event-related fMRI paradigm (Wakeman and Henson, 2015). Experiment 2 measured the sustained response to 14s blocks containing rapid presentations of oriented visual gratings, with the orientation alternating between 45 and 135 degrees across blocks (Alink et al., 2013).

**Figure 2.**
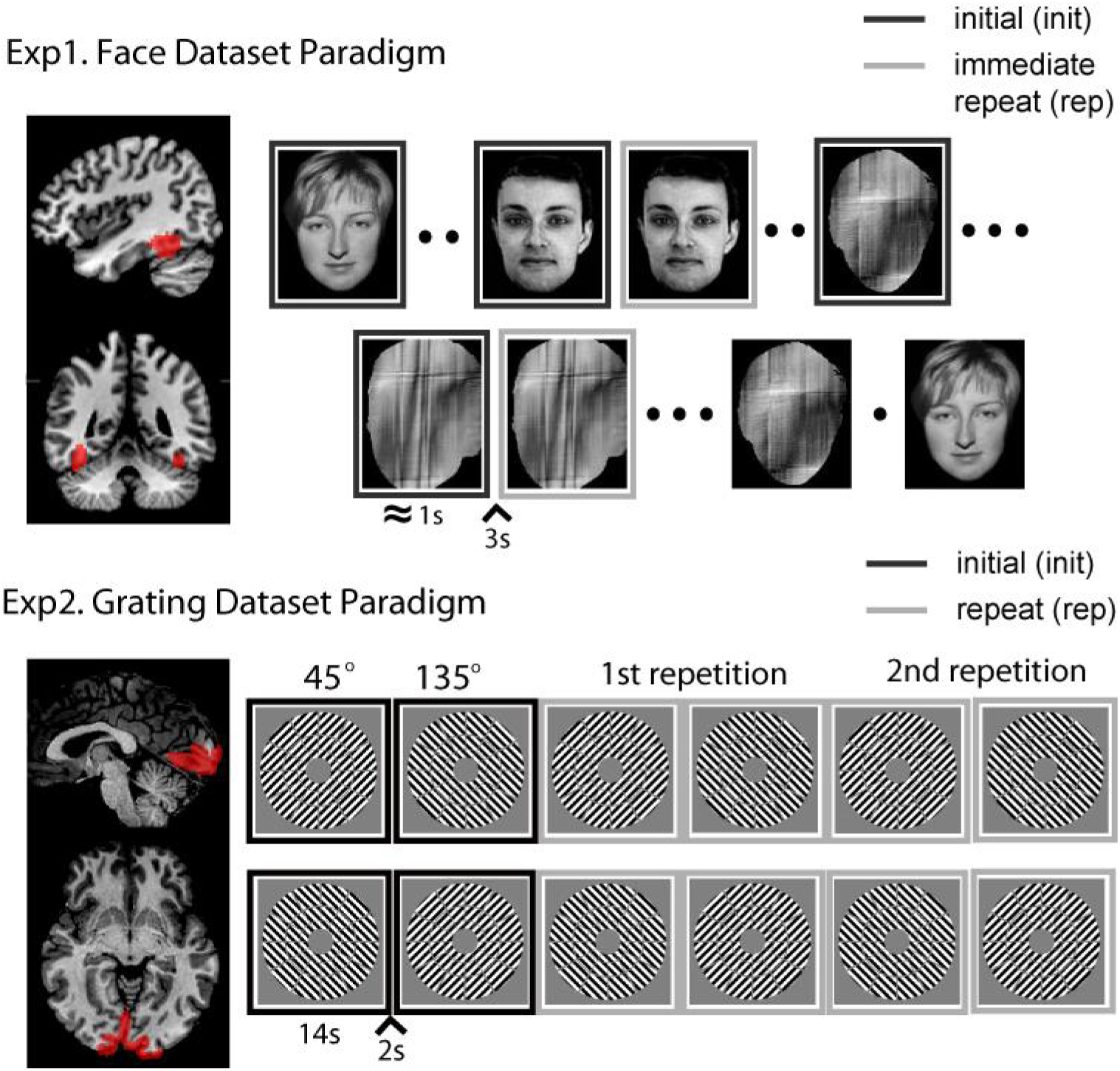
visualization of the two experimental paradigms and the corresponding brain region used in the analyses. Note in our analyses we randomly dropped half of the initial trials in the Face dataset to balance the number of the initial and immediate repeats.

The six data features are shown for each experiment in Figure 3. As expected, both Experiment 1 and Experiment 2 showed a significant repetition suppression (**MAM**) in FFA (t(17)=−7.53, P<.001) and V1 (t(17)=−7.13, P<.001), respectively. Stimulus repetition also reduced both within-class (**WC**) and between-class (**BC**) correlations between trials in both experiments (Exp1/FFA, WC, t(17)=−8.61, P<.001, and BC, t(17)=−5.84, P<.001; Exp2/V1, WC, t(17)=−7.19, P<.001, and BC, t(17)=−7.76, P<.001). However, while the difference between repetition effects on within- and between-class correlations (**CP**) increased for V1 in Experiment 2 (t(17)=+4.15, P<.001), it decreased for FFA in Experiment 1 (t(17)=−3.84, P=0.0012). In other words, repetition improved the ability to classify stimuli according to their two classes in Experiment 2 (consistent with the improved classification for predictable stimuli reported in Kok et al (Kok et al., 2012)), but impaired such classification in Experiment 1 (as confirmed by support-vector machines in both cases). Furthermore, while linear regression showed that repetition suppression decreased with mean amplitude (**AMA**) in both experiments (Exp1/FFA, t(17)=+9.26, P<.001; Exp2/V1, t(17)=+7.83, P<.001), its dependence on voxel selectivity (**AMS**) differed across experiments: increasing with selectivity in Experiment 1 (FFA, t(17)=+3.46, p=0.003), but decreasing with selectivity in Experiment 2 (V1, t(17)=−2.31 p=0.034). Thus, given their different ROIs, stimulus-types and stimulation protocols, it is interesting to note that there are both commonalities and differences in the effects of repetition across the two experiments.

**Figure 3.**
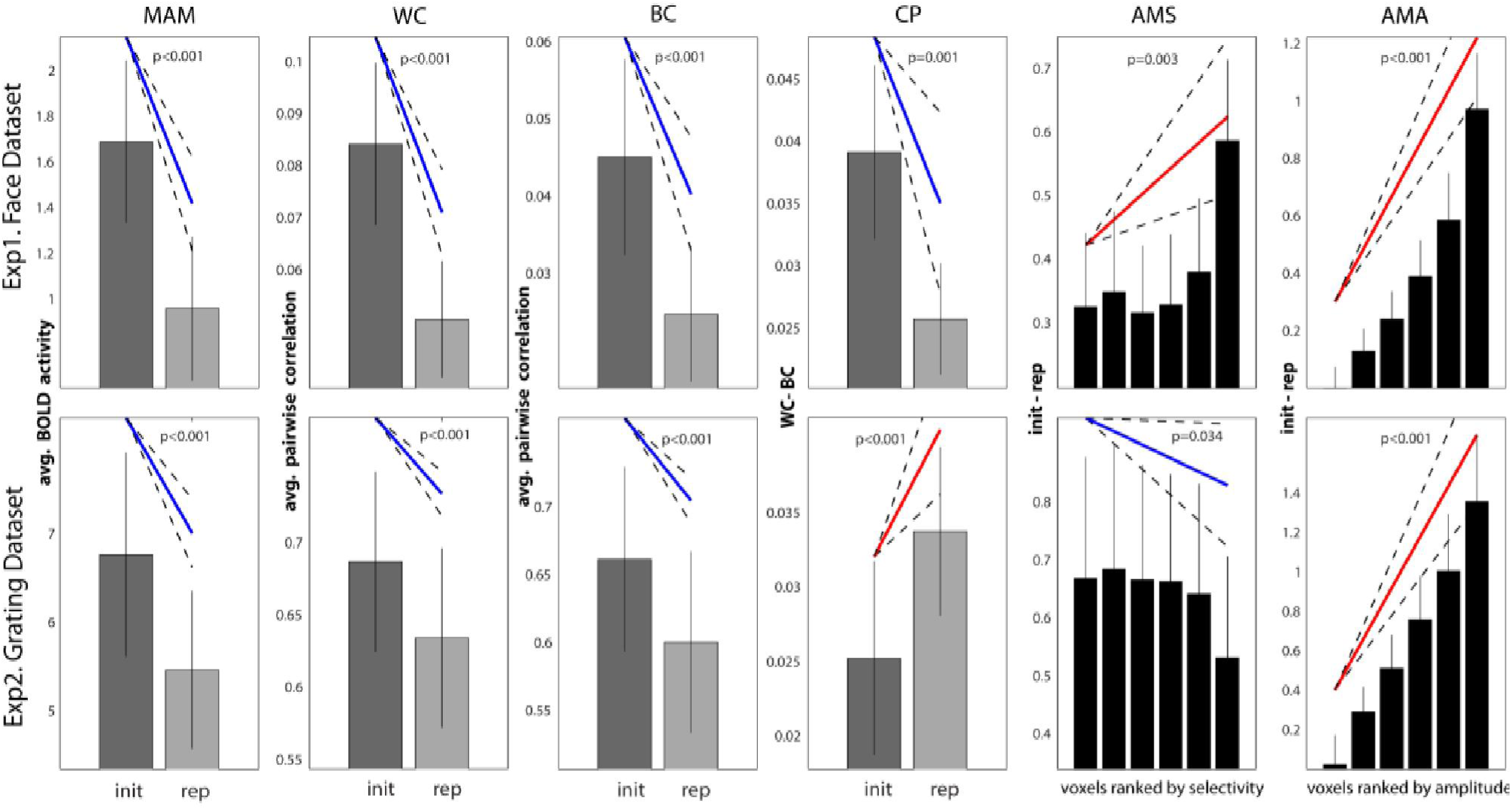
The six data features (columns) for each paradigm (rows). Bars reflect mean of each condition (init = initial presentation; rep = repeated presentation), with error bars reflecting 95% confidence interval given between-participant variability; diagonal line represents slope of linear contrast across conditions (red = positive; blue = negative) with dashed error margins reflecting 95% confidence interval of that slope (equivalent to pairwise difference when only two conditions).

### Simulation Results

We explored the predictions of each of the twelve models for each of the six data features in a grid search covering a wide range of values for the three free parameters (see Methods for specific ranges used). We ran 50 simulations for each model for all unique parameter combinations. For each such combination, we calculated the 99% confidence interval across the 50 simulations for the mean of each of the six data features, and tested whether this was above, below or overlapped zero.

#### Unconstrained parameters

In this initial analysis, different parameter values were allowed for each data feature, in order to see whether any data features were diagnostic of a neural model, and whether each model could, in principle, explain each data feature. Figure 4 summarizes the results for each experiment. The colors in each circle summarize the possible effects of repetition, i.e., whether various parameter combinations could produce an increase, decrease, no effect, or some combination of these.

**Figure 4.**
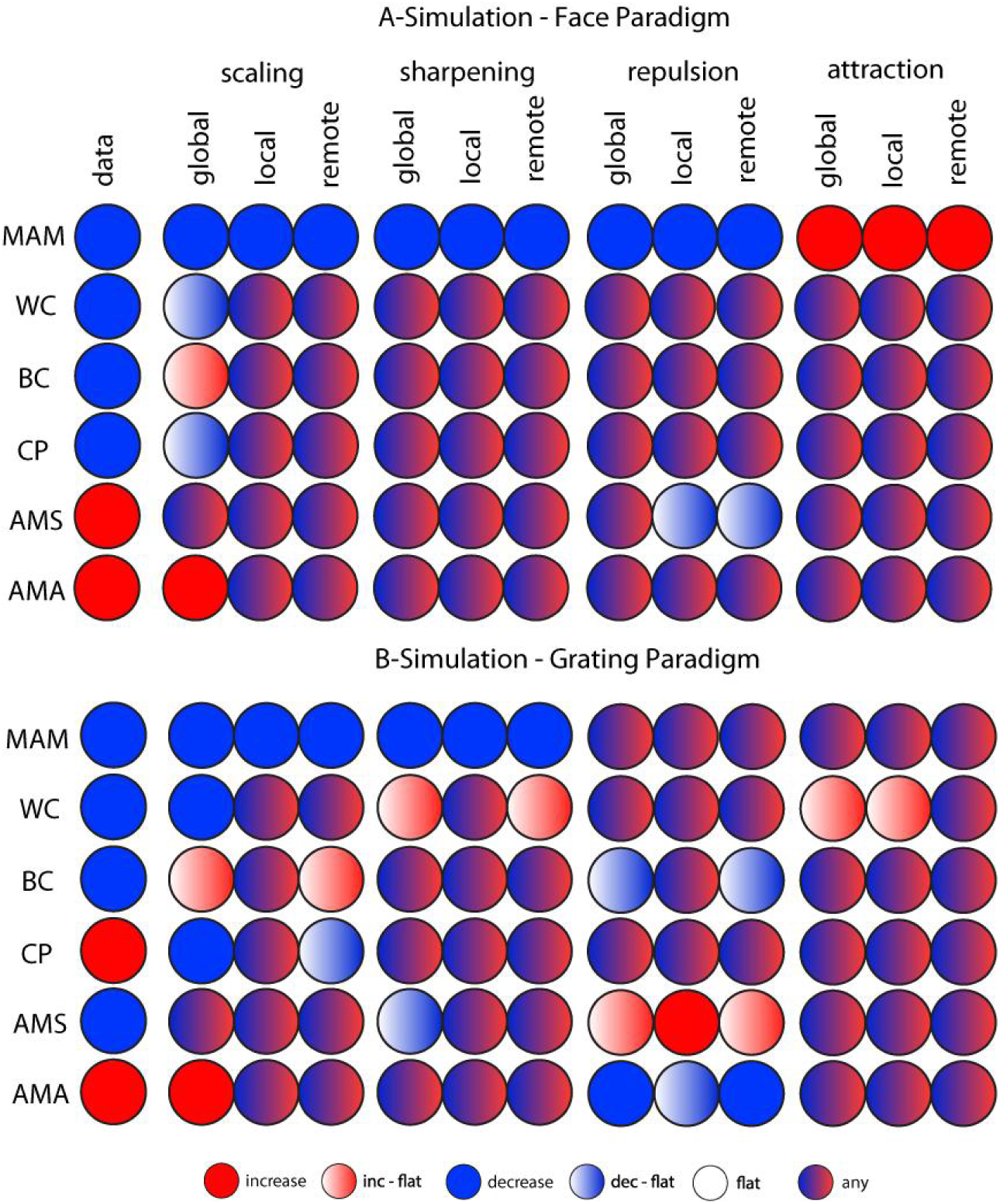
Possible simulated fMRI data features for all models (columns 2-13) for both experimental paradigms when considering all parameter combinations. The first column shows the empirical data features.

The first thing to note is that no single data feature was sufficient to identify the underlying neural model, illustrating the difficulty of inferring from fMRI data at the level of voxels to mechanisms at the level of neurons (i.e., no value in any row in Figure 4 is unique to one of the twelve models). Note that this conclusion holds regardless of the empirical value of the data features observed in the present experiments (leftmost column). This conclusion is important because some of these features, such as the increase in classification performance (CP) after repetition, have been assumed to support sharpening models (Kok et al., 2012), yet Figure 4 shows that several other non-sharpening models can produce an increase in CP. The same goes with the negative slope in AMS, which was also thought to support sharpening models (Kok et al., 2012).

The second thing to note is that some of the neural models cannot produce at least one of the data features observed in the present experiments (whatever their parameter settings within the large range explored here). This can be seen by comparing the leftmost column with the remaining twelve columns. This means that, by considering a range of consequences of repetition (both univariate and multivariate), one can at least rule out some neural models. Nonetheless, with unconstrained parameters, there were six models that could fit the data features in Experiment 1 (FFA) - local and remote scaling, all three sharpening models and global repulsion - and three models that could fit the data features in Experiment 2 (V1) - local scaling, local sharpening and remote attraction.

## Constrained parameters

Figure 4 shows results based on parameters whose values were varied for each data feature separately (e.g., the values *σ*, *a* and *b* that produce a decrease in MAM may not be the same values that produces a decrease in WC). When we constrained the parameters to have the same values across all six data features, only one model could simultaneously fit all six features: local scaling. This is shown in Figure 5, for which green and red colors now show whether a model could fit the data feature observed empirically (when parameter values were selected that explained the maximum number of features): only the column for the local scaling model has green colors for all data features. Moreover, this was the same model for both Experiment 1 and Experiment 2. The range of parameter values required for local scaling can be found in the Supplementary Table, and an illustrative fit to the data features can be found in Supplementary Figure 4 and the Supplementary Table. Therefore, the constraints offered by simultaneously fitting six different data features are sufficient to single out one of the twelve models considered here, and this was the same model across two independent experiments (just with different parameter values that are needed to capture differences between representations of the different stimuli across different ROIs, different stimulation protocols, etc.).

**Figure 5.**
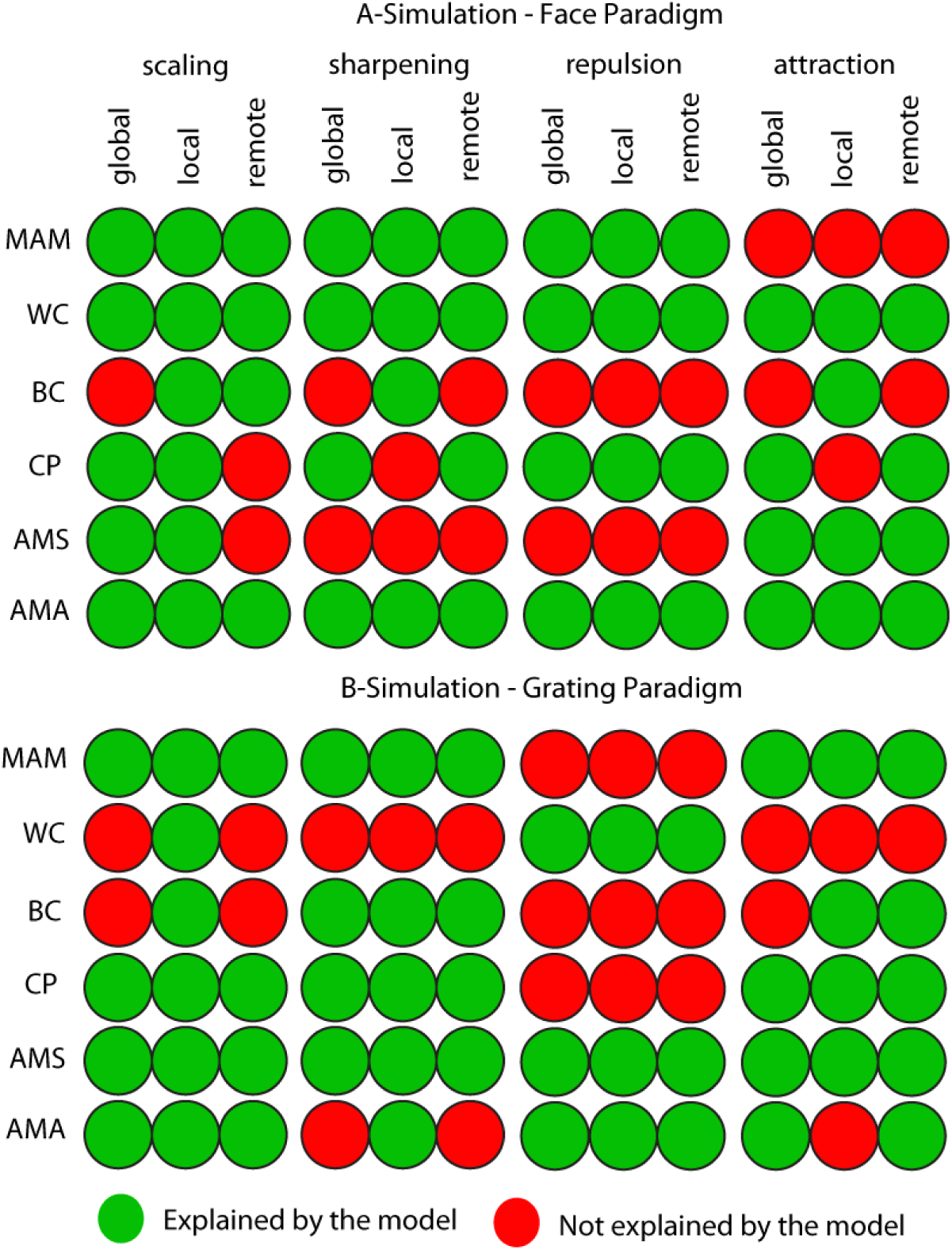
The maximum number of data properties explained by each model when parameters are constrained to be equal across all data properties (but can differ across experiments). Note that, for some models that can explain only 4 or 5 data properties with the same parameter values, there may be different subsets of the same number of data properties that can be explained (i.e., this figure only shows one such subset).

## Discussion

Several neural mechanisms have been hypothesized to underlie the observation of reduced fMRI responses to repeated stimulation, i.e. repetition suppression (Grill-Spector et al., 2006; Henson and Rugg, 2003; Naccache and Dehaene, 2001; Wiggs and Martin, 1998). These include neural habitation or fatigue, which down-scale neuronal firing rates, and neural tuning or sharpening, which tighten neuronal tuning curves (Grill-Spector et al., 2006). In principle, any of these mechanisms can explain the basic effect of repetition on the mean univariate fMRI response across voxels. However, we show that by 1) considering a range of features of fMRI repetition effects, both univariate and multivariate, and 2) formally modelling a range of potential neural mechanisms, various hypothetical neural mechanisms can be distinguished. Indeed, our results show that local scaling of neuronal firing is the only model, of the twelve considered here, that can simultaneously explain six features of repetition in fMRI data from two independent experiments. Importantly, local scaling at the neuronal level can explain sharpening of patterns at the voxel level (e.g. leading to improved classification after repetition), i.e., conceptually, sharpening at the voxel level does not imply sharpening at the neural level. This emphasizes the importance of formal modelling to bridge the differences between the neuronal and fMRI levels of investigation.

Our modelling also allowed us to demonstrate that no single feature of fMRI repetition effects was diagnostic of any of our twelve models (allowing for the 2-3 degrees of freedom corresponding to the parameters of each model). This reinforces the importance of simultaneously considering multiple data features. The local scaling model was the only model capable of simultaneously fitting our six data features with the same set of parameter values. This was true across datasets that differed in terms of the stimuli, brain region of interest, stimulation protocol and analysis method (randomly intermixed events versus sustained blocks of repetition). It is possible that these findings could be explained by combinations of mechanisms (e.g. global scaling and local sharpening), or by neuronal mechanisms beyond the twelve considered here. Nonetheless, local scaling remains the most likely current explanation, in terms of parsimony.

A study by Weiner and colleagues (Weiner et al., 2010) also used formal neural models to examine univariate fMRI repetition suppression (what we call here MAM). They assessed the validity of two models – comparable to our global scaling and global sharpening models – and found that the validity of each model depended both on brain area and lag between stimulus repetitions. Neither global scaling nor global sharpening however is able to account for the full range of repetition effects in our two fMRI datasets. Nonetheless, Weiner et al.’s investigations raise the important possibility that the neural mechanisms underlying fMRI repetition effects may vary with other factors, e.g., longer lags may engage qualitatively different neural mechanisms than the immediate repetition examined here (Ashida et al., 2007; Cavina-Pratesi et al., 2010; Chong et al., 2008; Fang et al., 2005; Fox et al., 2009; Grill-Spector et al., 2006; Henson, 2016b; Kar and Krekelberg, 2016; Krekelberg et al., 2006; Winston et al., 2004).

To our knowledge, only Kok and colleagues (Kok et al., 2012) have previously examined a combination of univariate and multivariate fMRI data features to elucidate neural mechanisms of fMRI response reductions. Specifically, they observed that stimulus predictability reduced mean fMRI responses (MAM) while increasing fMRI pattern information about stimulus class (what we call here CP). This observation, in conjunction with their finding that fMRI response reduction was reduced for voxels with higher stimulus selectivity (what we call here AMS), was interpreted as evidence for (global) neural sharpening. For the grating experiment (Experiment 2), we observed a similar pattern of empirical results. Although we did not explicitly manipulate prediction, the clear block structure of repeating stimuli would have produced strong predictions for upcoming stimuli (the same was not true for the face experiment, which may be why it showed the opposite pattern for AMS). Importantly however, our simulations revealed that this conjunction of data features does not, in fact, uniquely identify neural sharpening. Nonetheless, our results do not directly contradict the sharpening claim of Kok and colleagues (Kok et al., 2012), in which the paradigm differs from that employed in the two current experiments; rather the models considered here would need to be applied to Kok et al’s data to see if local scaling was again the only model able to explain the full pattern of results, or whether local sharpening was better in this case.

Our finding that local scaling best explains fMRI repetition suppression does not question previous findings of stimulus repetition effects on single-cell recordings (Bachatene et al., 2015; Sur et al., 2001; Jeyabalaratnam et al., 2013; Kar and Krekelberg, 2016; Ringach and Bredfeldt, 2002; Swindale, 1998). As alluded to above, it is possible that multiple mechanisms operate in parallel, but in different neural populations or cortical layers, and that the dominance of the local scaling model is simply due the greatest proportion of neurons exhibiting local scaling. It is also important to keep in mind that our fMRI signals are dominated by changes in local field potentials (Goense and Logothetis, 2008), rather than the action potentials in large pyramidal cells that are normally measured in single-cell studies.

Note that local scaling of neuronal tuning curves could itself arise from multiple potential mechanisms within the context of a neuronal circuit, such as synaptic depression of bottom-up inputs, or recurrent inhibition by top-down inputs. For example, the hypothesis of predictive coding, which has been used to explain other aspects of repetition suppression (Henson and Rugg, 2003; Kiebel et al., 2009), would also result in maximal suppression of neurons that are most selective for the (repeated) stimulus – i.e., local scaling. Moreover, our fMRI data cannot speak to the temporal dynamics of repetition effects, such as the acceleration of evoked responses predicted by facilitation models (Henson, 2016a) and neural synchronization (Gotts et al., 2012).

In sum, our work illustrates the value using forward models that map from neurons to voxels, like those considered here, to interpret fMRI data, i.e., map back from voxels to neurons. Despite the simplicity of the models considered here, their predictions are not always intuitive, and they therefore help protect against superficial analogies, for example that sharpening of multivoxel fMRI patterns entails the sharpening of neuronal tuning curves.

## Methods

### Experimental Details

#### Experiment 1 – Face data

##### Participants and Task

We report data from 18 of the 19 participants (8 female, aged 23-37) described in Wakeman & Henson (Wakeman and Henson, 2015), after removing one participant who had fewer fMRI volumes than the others. In brief, participants made left-right symmetry judgments to faces or scrambled faces. Half of the faces were famous, but only the remaining unfamiliar faces were analysed here. Every stimulus was repeated either immediately or after a delay of 5-15 intervening stimuli; only initial presentations and immediate repetitions are analysed here. For more details, see Wakeman & Henson (Wakeman and Henson, 2015).

##### fMRI acquisition & processing

The MRI data were acquired with a 3T Siemens Tim-Trio MRI scanner with 32-channel headcoil (Siemens, Erlangen, Germany). The functional data were acquired using an EPI sequence of 33, 3 mm-thick axial slices (TR 2000 ms, TE 30 ms, flip angle 78°). 210 volumes were acquired for each of 9 sessions. Slices were acquired in an interleaved fashion, with odd then even numbered slices and a distance factor adjusted to ensure whole-brain coverage, resulting in a range of native voxel sizes of 3×3×3.75 mm to 3×3×4.05 mm across participants (for details see (Wakeman and Henson, 2015)). We also obtained a high-resolution (1mm isotropic) T1-weighted anatomical image using a Siemens MPRAGE sequence for normalisation of brains.

The fMRI data were preprocessed using the SPM12 software package (www.fil.ion.ucl.ac.uk/spm) in Matlab 2012b (uk.mathworks.com). After removing the first two EPI images from each session to allow for T1 saturation effects, the functional data were corrected for the different slice times, realigned to correct for head motion, and coregistered with the structural image. The structural image was warped to a standard template image in MNI space, and the warps then applied to the functional data.

We extracted fMRI timeseries data from voxels within a combined left and right Fusiform Face Area (FFA) mask, as defined by the group univariate contrast of unfamiliar faces versus scrambled faces (averaged across initial and repeated presentations), thresholded at p<.05 family-wise error corrected using random field theory. The combined masked contained 135 voxels for the right FFA and 50 voxels for the left FFA.

Since this design used a short SOA of between 2.9-3.3s (randomly jittered), the BOLD response for each trial was estimated using the Least Squares Separate (LSS-N) approach (Mumford et al., 2012), (Abdulrahman and Henson, 2016), where N is the number of conditions (qualitatively similar results were achieved with the standard Least Squares All approach). LSS-N fits a separate General Linear Model (GLM) for each trial, with one regressor for the trial of interest, and one regressor for all other trials of each condition (plus 6 regressors for the movement parameters from image realignment, to capture residual movement artefacts). This implements a form of temporal smoothness regularisation on the parameter estimation (Abdulrahman and Henson, 2016). The regressors were created by convolving a delta function at the onset of each stimulus with a canonical haemodynamic response function (HRF). The parameters for the regressor of interest for each voxel in the mask were then estimated using ordinary least squares, and the whole process repeated for each separate trial. The number of trials varied slightly from session to session but it was balanced across participants, totalling 49 trials for each of the 4 trial-types considered here (initial and immediate repetitions of unfamiliar and scrambled faces).

#### Experiment 2 – Visual grating dataset

##### Participants and Task

We report fMRI data from the visual gratings conditions of a larger study reported previously (Alink et al., 2015; 2013; 2016). Eighteen healthy volunteers (13 female, age range 20–39) with normal or corrected-to-normal vision took part in the experiment.

The gratings were oriented 45° clockwise and 45° anticlockwise (135° clockwise) from the vertical, with a spatial frequency of 1.25 cycles per visual degree (a frequency that strongly drives V1 (Henriksson et al., 2008)). These stimuli were presented during 2 runs of 8 minutes, with each run divided into 4 subruns, and each subrun containing 6 blocks, with the orientation presented in each block alternating (Figure 1). Each block lasted 14s and contained 28 phase-randomized gratings of one orientation, presented at a frequency of 2 Hz. The stimulus duration was 250 ms, followed by an interstimulus interval (ISI) of 250 ms, during which a central dot was present, surrounded by a ring that determined the task (see below). The spatial phase was drawn randomly from a uniform distribution between 0 and 2π. Stimulus blocks were separated by 2s fixation periods and subruns by 24s fixation periods.

In addition, each participant participated in a 12-minute run for retinotopic mapping. A description of the stimuli employed and the procedure used to define individual regions of interest (ROIs) for the primary visual cortex can be found in Alink et al. (Alink et al., 2013).

Participants were instructed to continuously fixate on a central dot (diameter: 0.06° visual angle). The dot was surrounded by a black ring (diameter: 0.20°, line width: 0.03°), which had a tiny gap (0.03°) either on the left or right side. The gap switched sides at random at an average rate of once per 3 s (with a minimum inter-switch time of 1s). The participant’s task was to continuously report the side of the gap by keeping the left button pressed with the right index finger whenever the gap was on the left side, and keeping the right button pressed with the right middle finger whenever the gap was on the right side. The task served to enforce fixation and to draw attention away from the stimuli.

##### fMRI acquisition & processing

Functional and anatomical MRI data were acquired on the same scanner as Experiment 1 (see above). During each stimulus run, we acquired 252 volumes containing 31 transverse slices covering the occipital lobe as well as inferior parietal, inferior frontal, and superior temporal regions for each subject using an EPI sequence (TR=2000ms, TE=30ms, flip angle=77°, voxel size: 2.0mm isotropic, field of view: 205mm; interleaved acquisition, GRAPPA acceleration factor: 2). The same EPI sequence was employed for a retinotopic mapping run, during which we acquired 360 volumes. We also obtained a high-resolution (1mm isotropic) T1-weighted anatomical image using a Siemens MPRAGE sequence.

Functional and anatomical MRI data were preprocessed using the Brainvoyager QX software package (Brain Innovation, v2.4). After discarding the first two EPI images for each run to allow for T1 saturation effects, the functional data were corrected for the different slice times and for head motion, detrended for linear drift, and temporally high-pass filtered to 2 cycles per run. The data were aligned with the anatomical image and transformed into Talairach space (Talairach and Tournoux, 1988). After automatic correction for spatial inhomogeneities of the anatomical image, we created an inflated cortex reconstruction for each subject. All ROIs were defined in each individual subject’s cortex reconstruction and projected back into voxel space.

For simplicity, we contrasted first versus third presentations of each orientation within a subrun, to maximize repetition effects. There are likely to be effects induced by intermediate presentations of the other orientation, which are modelled fully in the models described below. Any effects of the order of specific orientations were controlled by counterbalancing (i.e., averaging data over 45°−135°−45°−135°−45°−135° and 135°−45°−135°−45°−135°−45° subruns).

### fMRI Data Properties

Let *B_vtpc_* be the BOLD signal at voxel *v* for trial *t* involving presentation *p* of stimulus class *c* (where *c* is face or scrambled face in Experiment 1, or 45 versus 135 degrees in Experiment 2). The number of voxels (*N_v_*) was 185 for the FFA ROI in Experiment 1, and varied from 775-1598 across participants (M=1100, SD=220) for the V1 ROI in Experiment 2. The number of trials (replications) for each stimulus class and presentation (*Nt*) corresponded to 49 events in Experiment 1, and the 8 sub-runs in Experiment 2. We identified 6 properties of the fMRI data, based on various analyses that have been reported in various fMRI studies (Kok et al., 2012; Weiner et al., 2010), though never investigated simultaneously within a single study:

**1-Mean Amplitude Modulation (MAM):** This represents the mean, over voxels, trials and the two classes, of the difference in the fMRI response to the initial versus repeated presentations:

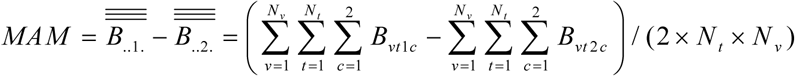

This is the typical univariate measure of repetition suppression (Grill-Spector et al., 2006).

**2-Within Class Correlations (WC):** This is the mean pairwise correlation of patterns over voxels, averaged across all trials and classes, and then contrasted for initial versus repeated presentations:

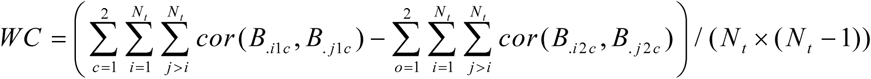

This captures how repetition makes patterns for the same class more or less similar.

**3-Between Class Correlations (BC):** This is the mean pairwise correlation of patterns over voxels for all trials of different classes, contrasted for initial versus repeated presentations:

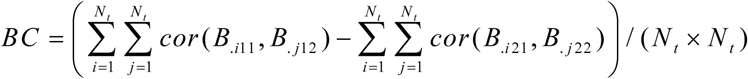

This captures how repetition makes patterns of the opposite class more or less similar.

**4-Classification Performance (CP):** The ability of MVPA to classify the two classes relates to the difference between Within- and Between-Class correlations:

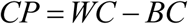

Note that this measure it is not redundant with the previous two features, since repetition might decrease both WC and BC, but decrease BC more, for example, such that CP increases.

To confirm the relationship between CP and MVPA classification, we compared our CP results to the results of an analysis using linear Support Vector Machine classification (SVM, Bioinformatics Toolbox in Matlab 2012b) with leave-one-session-out cross-validation for the face dataset and with leave-one-subrun-out cross-validation for the grating dataset. Classification accuracy was found to be higher for initial than for repeated presentations of face stimuli (initial: 70%, repeated: 61%, t(17) = 3,06, p < .005) while the opposite was observed for grating stimuli (initial: 56%, repeated: 69%, t(17) = −3,09, p < .005) – which fits with our our CP results.

**5-Amplitude Modulation by Selectivity (AMS):** This is a further breakdown of the first feature above, where the degree of amplitude modulation is related to the degree of “selectivity” of each voxel. Thus voxels were first binned (into six bins) by the absolute t-value of the difference between mean activity to each class (averaging across trials and both presentations, to avoid regression-to-the-mean), and then the slope estimated of a linear regression of repetition-related modulation against selectivity bin:

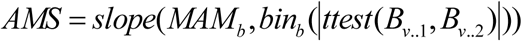

where *slope*() is the slope of best-fitting linear function, *bin_b_*(*t*) bins voxels to the six bins according to ascending values of *t* from a *t*-test at each voxel across all trials of each condition, and *MAM_b_* is the amplitude of the repetition effect averaged across all voxels in bin *b*. A negative slope indicates that adaptation suppresses non-selective voxels more than selective ones.

**6-Amplitude Modulation by Amplitude (AMA):** This is identical to AMS above, except that voxels were binned by amplitude (averaging across trials, classes and presentations) rather than by selectivity.

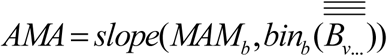

A positive slope means that adaptation suppresses more responsive voxels more, which Weiner et al (Weiner et al., 2010) claimed is indicative of scaling models.

#### Modelling approach

Assuming that neural populations (e.g. orientation columns) have a unimodal tuning-curve along a relevant stimulus dimension (e.g., orientation), at least four basic neural mechanisms of adaptation have been suggested: 1) scaling, where neural populations reduce their firing rate, in proportion to their initial firing rate, i.e, their tuning curves are suppressed (Kar and Krekelberg, 2016) (also called neural “fatigue” (Grill-Spector et al., 2006; Ringach and Bredfeldt, 2002; Swindale, 1998)), 2) sharpening, where the width of neural tuning curves decreases (Kar and Krekelberg, 2016), 3) repulsive shifting, where the peaks of tuning curves shift away from the adaptor (Dragoi et al., 2001) and 4) attractive shifting, where the peaks shift towards the adaptor (Jeyabalaratnam et al., 2013). These four mechanisms can be further parametrized according to whether the adaptation is 1) global, affecting all neural populations regardless of their preferred stimulus, 2) local, where adaptation is greater for neural populations whose preferences are closer to the adaptor, and 3) remote, where adaptation is greater for neural populations whose preferences are further from the adaptor. This results in a space of twelve possible models, as defined formally below.

##### Simulating neural responses

For simplicity, neuronal tuning curves were assumed to lie on a single stimulus dimension. These tuning curves where characterised by the firing rate, *f_i_*(*j*), for the *i*-th neural population in response to (the first presentation of) stimulus *j*:

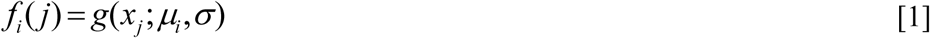

where *g*(*x_j_*;*μ_i_*,*σ*) is a function of the value of stimulus *j*, *x_j_*, parametrised by the preference of the *i*-th neural population (peak of its tuning curve), *μ_i_*, and the width of the tuning curve, *σ*. The value of *x_j_* was bounded from 0 *X*, and for comparability with the circular dimension of orientation for Experiment 2 (see below), *X* = *π*. The preferred stimulus for each neural population (*μ_i_*) was selected randomly from a uniform distribution across this range, whereas the tuning width was assumed equal for all populations.

For Experiment 1, the single dimension represented faces, and the tuning curve *g*(*θ_j_*; *μ_i_*, *σ*) was a Gaussian function (such that *μ_i_* was its mean and *σ* was its standard deviation). For Experiment 2, the stimulus dimension represented the orientation of gratings, which is a circular dimension, and so tuning curves were modelled by a von Mises distribution (Ringach and Bredfeldt, 2002; Swindale, 1998). Given that the response of orientation columns is symmetrical around 180^o^, *x_j_* varied from 0 *π* radians, such that the von Mises distribution was defined as *g*(*θ_j_*; *μ_i_*, *σ*)=*VM*(2*x_j_*;2*μ_i_*,1/*σ*). Both distributions were normalised to have a peak height of 1 (i.e, the tuning curves do not represent probability distributions).

As in Weiner et al (Weiner et al., 2010), the extent of repetition suppression was expressed through the variable *0*<*c*<*1*. Thus according to the four basic neural mechanisms of adaptation, the firing rate in response to second presentation of stimulus *j* was:

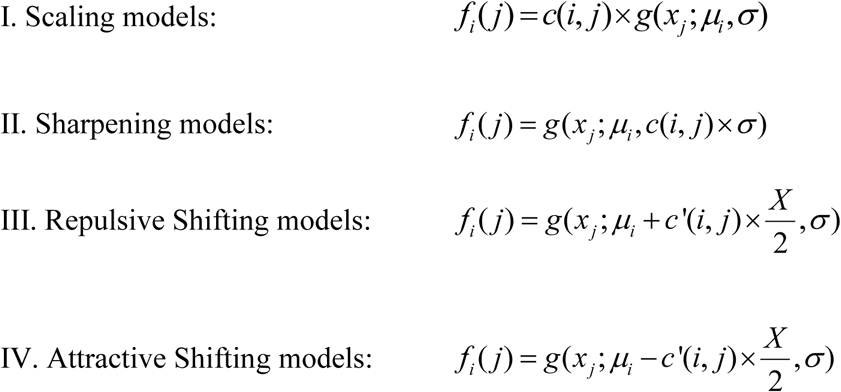

where *c*′(*i*, *j*), for shifting models, is defined below, and where *g*, for all models except scaling, is a re-normalised version of *g* such that its peak height remains 1 (i.e, we assumed that sharpening and shifting do not affect the peak firing rate).

Unlike Weiner et al. (Weiner et al., 2010), *c* was itself a function of the distance between the neural preference and stimulus value, i.e, *c*(*i*, *j*) = *h*(*d*(*i*, *j*);*a*,*b*) where *d*(*i*, *j*) = *μ_i_* – *x_j_* and *a* and *b* are free parameters that control the domain over which neural adaptation applied. (Note that, for the circular dimension in Experiment 2, the distance function is also circularised to *d* (*i*, *j*) = min (|*d* (*i*, *j*)|,2*X* – |*d*(*i*, *j*)|)). The parameter 0<*a*<*1* controlled the maximal adaptation, while 0<*b*<*X*/*2* controlled how rapidly adaptation changed with the distance between neural preference and stimulus property. Three different distance functions were considered:

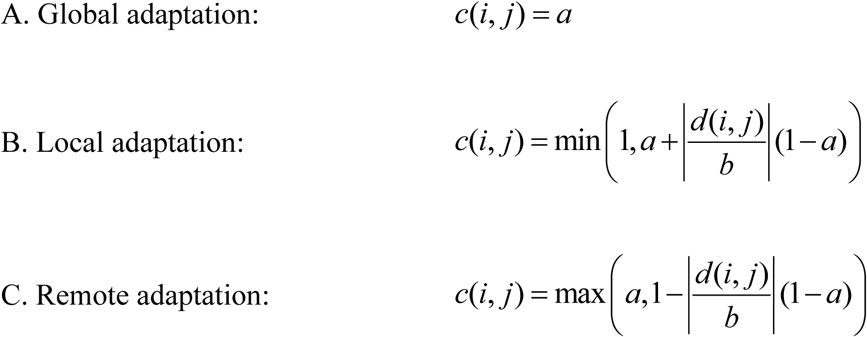

The parameter *b* represents a linear slope, and in combination with the min/max operations, provides a piecewise linear function that implements the simplest form of a nonlinear saturation, as shown in Supplementary Figure 2. This nonlinearity is important for Experiment 2, to break the symmetry of results after adapting to two opposite orientations (otherwise the response of neural populations whose preference is half-way between the two adaptors could never exceed that of neural populations whose preference matches either adaptor). (Global adaptation is a special case of Remote adaptation when *b* = 0, and a special case of Local adaptation when *b* = ∞.)

Finally, for the two shifting models, the adaptation factor *c*′(*i*, *j*) additionally 1) depended on the sign of the difference between neural preference and stimulus value:, i.e., *c*′(*i*, *j*) = sign(*d*(*i*, *j*)) × *c*(*i*, *j*), and 2) was defined such that *c*′(*i* = *j*) = 0, which means that even for local shifting, the population whose preferred stimulus matches the adaptator is not affected by adaptation. This is what is found empirically (Dragoi et al., 2001; 2000) and would actually correspond to a quadratic distance function. Rather than parametrize this quadratic function further, by defining *c*′(*i* = *j*) = 0, we are effectively limiting this quadratic function to the level of discretization of stimulus angles (*π*/8 here).

For multiple presentations of stimuli, we simply multiply the effects of *c*. Thus for the *p*-th presentation of a stimulus (*p*>1):

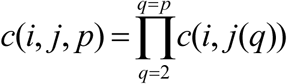

where *j*(*q*) represents the sequence of stimuli presented. For Experiment 1, each stimulus was only repeated once, so *p*=2. For Experiment 2, the two types of stimuli alternated across 6 blocks, such that *j* = [*π*/4,3*π*/4,*π*/4,3*π*/4,*π*/4,3*π*/4] or *j* = [3*π*/4,*π*/4,3*π*/4,*π*/4,3*π*/4,*π*/4] (with the initial stimulus type counterbalanced across subruns). Given the gap between subruns, we assumed adaptation wore-off between subruns, by resetting *f* to Equation 1 (which is supported by the activity patterns across the whole experiment shown in Supplementary Figure 3). Note also that the pattern of simulation results below remained unchanged if we additionally simulated repetition effects occurring within each block (i.e., affecting mean response to the first block too).

###### Simulating voxel responses

Each voxel was assumed to contain *N* neural populations, whose preferences were randomly selected from a uniform distribution (see below). Since the BOLD response is proportional to the neural firing rate (Heeger et al., 2000; Rees et al., 2000), the voxel response was simply the average firing rate of each population within that voxel. The number of neural populations per voxel, *N*, does not have a qualitative effect on the simulation results. However, it does have a quantitative effect: When *N* is large, the majority of the voxels would be similar to each other in their response, with very weak overall voxel biases towards particular stimulus classes. If *N* is small however, the voxels have stronger biases, and the quantitative differences among the models become more evident. Here we used *N*=8.

We then simulated *V*=200 voxels, and added a small amount of independent noise to each voxel, drawn from a Gaussian distribution with standard deviation of 0.1 (which represented an amplitude SNR of 10, since the peak value of *f_i_*(*j*) to initial presentations was scaled to 1).

To generate voxels that vary in their selectivity and activity, the value of *μ_i_* was sampled randomly with uniform probability from 8 possible orientations from *θ* = *0*…*7π*/*8* in steps of *π*/8. For Experiment 2, these values therefore included two neural populations that responded optimally to one of the stimuli (tuned to orientations 45^0^ and 135^0^), two highly non-selective neurons (tuned to orientations 0^0^ and 90^0^), and four partially-selective neurons in between. This allowed us to generate a sufficient variety of voxel activity and types, ranging from highly-selective voxels to partially-selective to non-selective voxels for each orientation.

Each model had 3 free parameters: *a*, *b* and *σ* (except for the Global models where there was no *b* parameter). We explored the predictions of each of the twelve models for each of the 6 data features in a grid search covering a wide range of values for the three parameters. The *a* values ranged from 0.1 to 0.9 in steps of 0.1 to cover a wide range of maximal adaptation, while *b* values ranged from 0.1 to π/2; in steps of 0.2 radians. For σ, values ranged from 0.1 to 1 in steps of 0.2, and then from 2 to twelve in steps of 3 to cover a wide range of tuning widths. For each mode, we ran 50 simulations for each of the 8000 unique combinations of these three parameters (or 800 for Global models with just two parameters). For each parameter combination, we calculated the 99% confidence interval across the 50 simulations for the value associated with each of the 6 data properties, and tested whether this was above, below or overlapped zero. Figures 4 and 5 summarise the results.

The Matlab code for these simulations (together with ROI data used in this paper) is available here: http://www.mrc-cbu.cam.ac.uk/people/rik.henson/personal/data4papers/

The fitted data properties for local scaling (using parameter values *a*=*0.8*, *b*=*0.4*, *s*=*0.4* for grating dataset and values a=0.7, b=0.2, s=0.2 for the face dataset) are shown in Supplementary Figure 4. Note that, while all the qualitative effects of repetition are reproduced successfully, the absolute values of BC correlations in our simulations are negative rather than positive.

The positive correlations of BC in the data could owe to several factors, such as correlated scanner noise or temporal drift (extrinsic scanner factors). Alternatively there could be intrinsic factors such as neural populations within a voxel that are not selective, responding equally to both stimulus classes (i.e flat tuning curves). Such diversity in the neural tuning curves has been reported in single cell literature (Schmolesky et al., 2000; Shapley et al., 2003). Hence, quantitative fitting of the data would require additional assumptions and scaling parameters that are not of theoretical interest in this paper. However, as a sanity check, we confirmed that we were able to achieve a positive BC correlation by adding a proportion of neural populations with flat tuning curves that respond and adapt equally to both stimulus types (Supplementary Figure 5). It is worth to mention that this addition did not change the overall conclusions of this paper, i.e. local scaling was still the wining model in both datasets even after adding this extra type of neurons to emulate correlated neural activity.

## Supplementary Material

**Sup. Figure 1:**
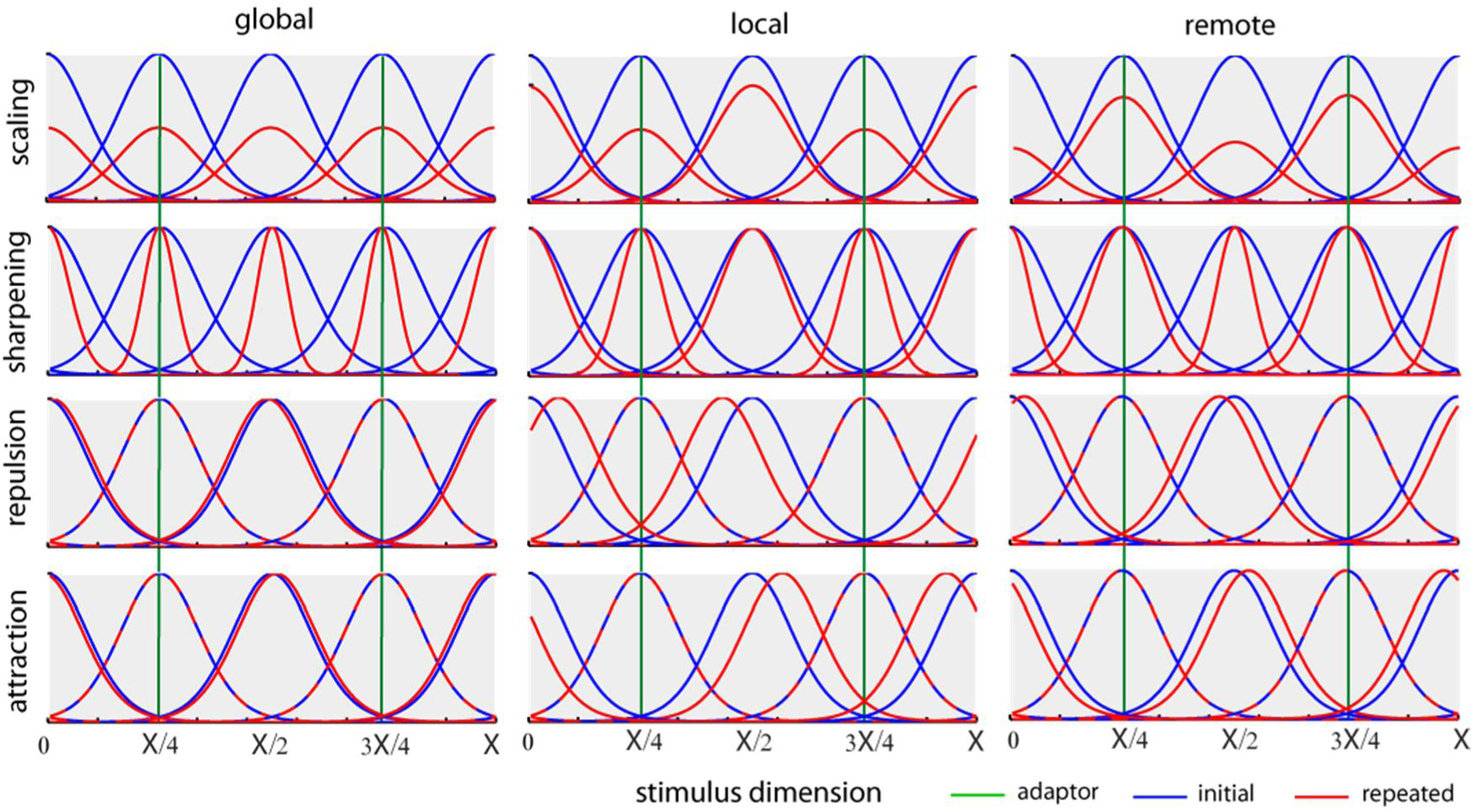
Example tuning curves before (blue) and after (red) after adaptation to both orientations in Experiment 2 according to the twelve different neural models of adaptation. Due to the circular nature of the stimulus dimension orientation, tuning curves were modelled with a von Mises distribution. For illustrative purposes, we only show four neural populations equally-spaced along the stimulus dimension.

**Sup. Figure 2:**
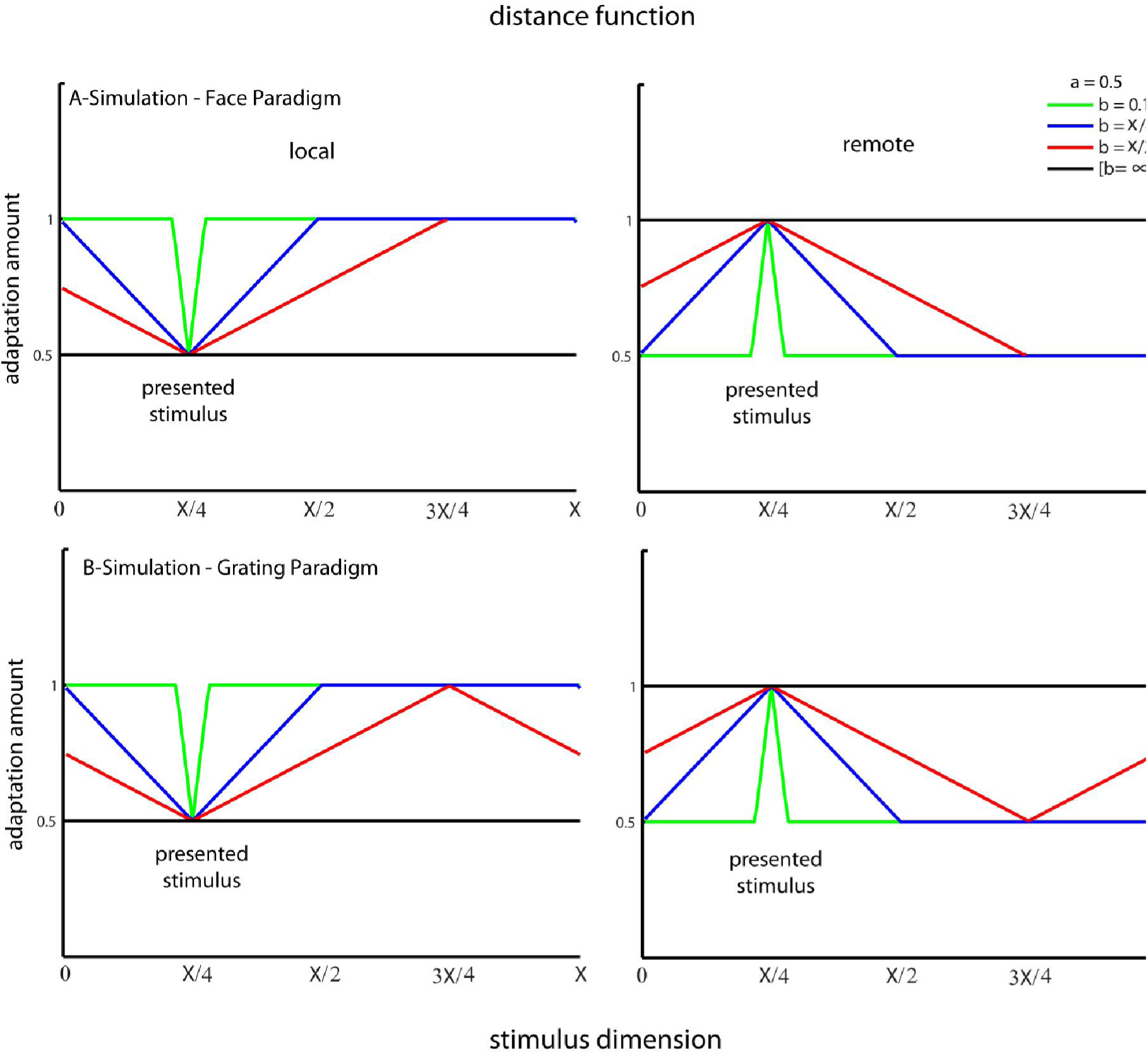
Distance functions, showing how amount of repetition suppression depends on distance between stimulus orientation (x-axis) and neural preference (here X/4) for a non-circular (top) and circular (bottom) dimension for local (left) and remote (right) domains. The *c* parameter is fixed to 0.5, while the *b* parameter is shown from 0.1 to ∞, though note that in our simulations, *b* only ranged from 0.1 to d=X/2.

**Sup. Fig3:**
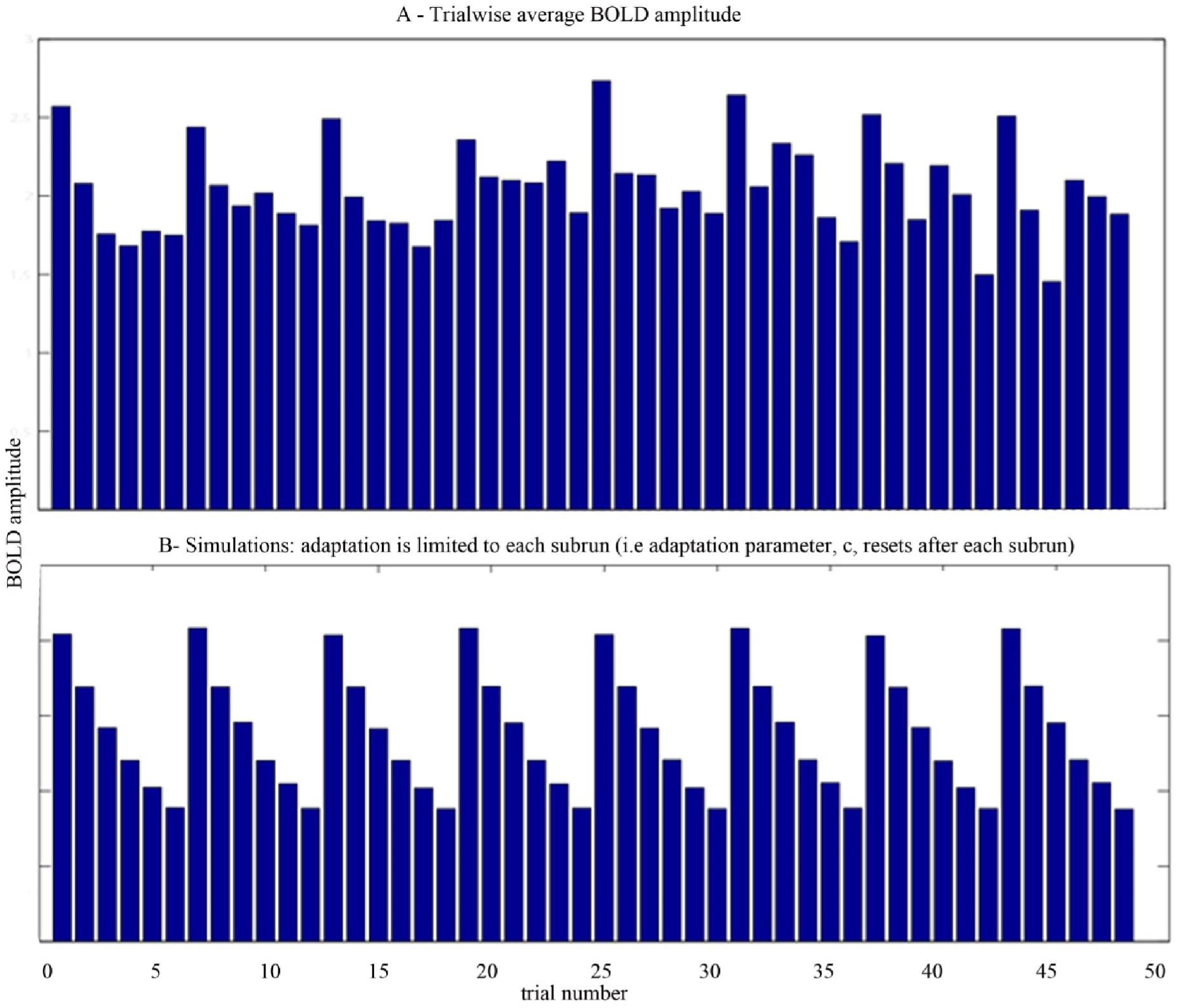
Showing A) adaptation effects were limited to the subruns and the BOLD activity at the start of each subrun were the same as the first trials in each subrun B) showing the effect of re-setting the adaptation factor between the independent runs in our simulation to match the empirical data results.

**Sup. Figure 4.**
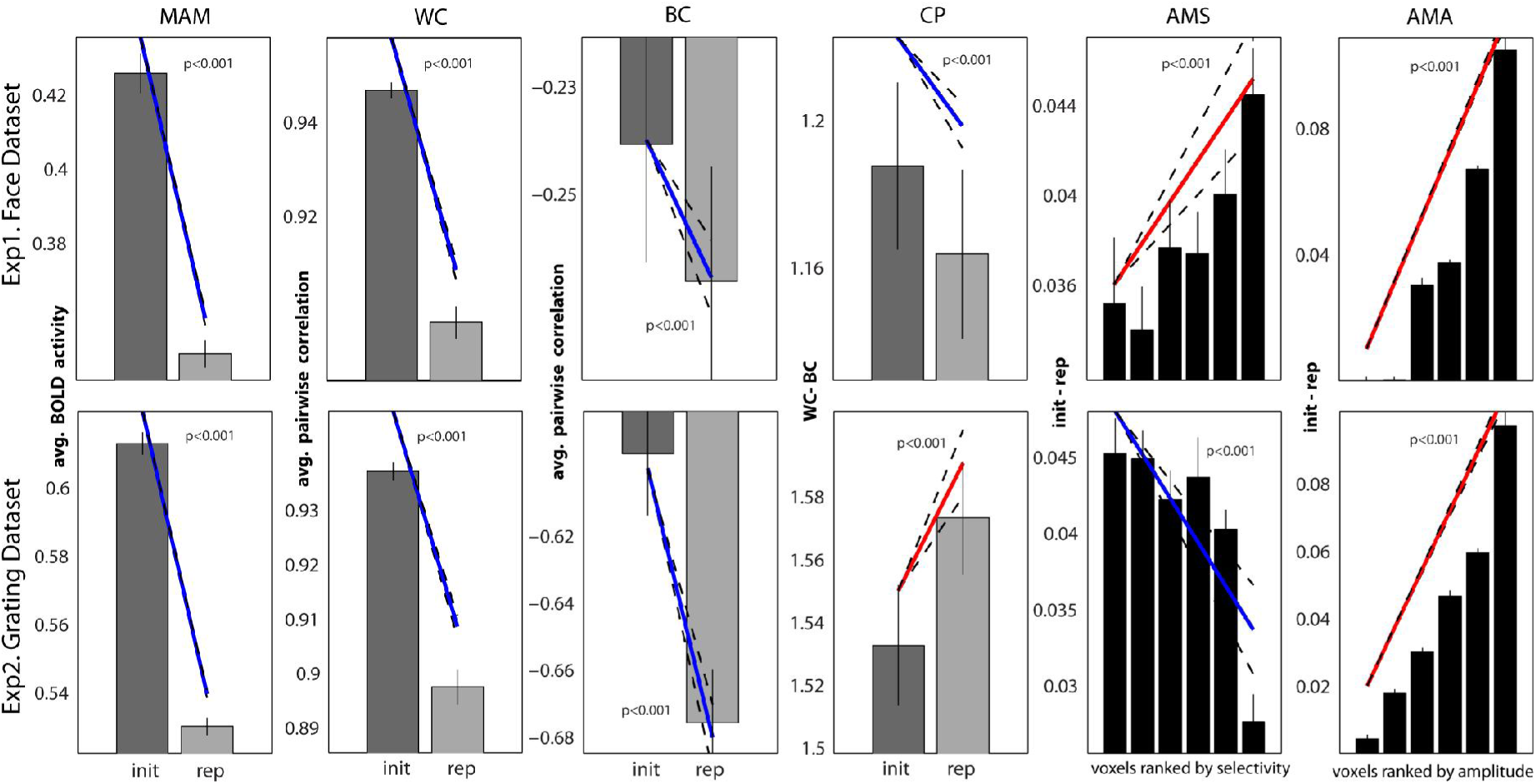
Local scaling prediction for all the 6 criteria with a sample wining parameter: c=0.8, b=0.4, sig=0.4 (grating dataset) and c=0.7, b=0.2, sig=0.2 (Face dataset)

**Sup. Figure 5:**
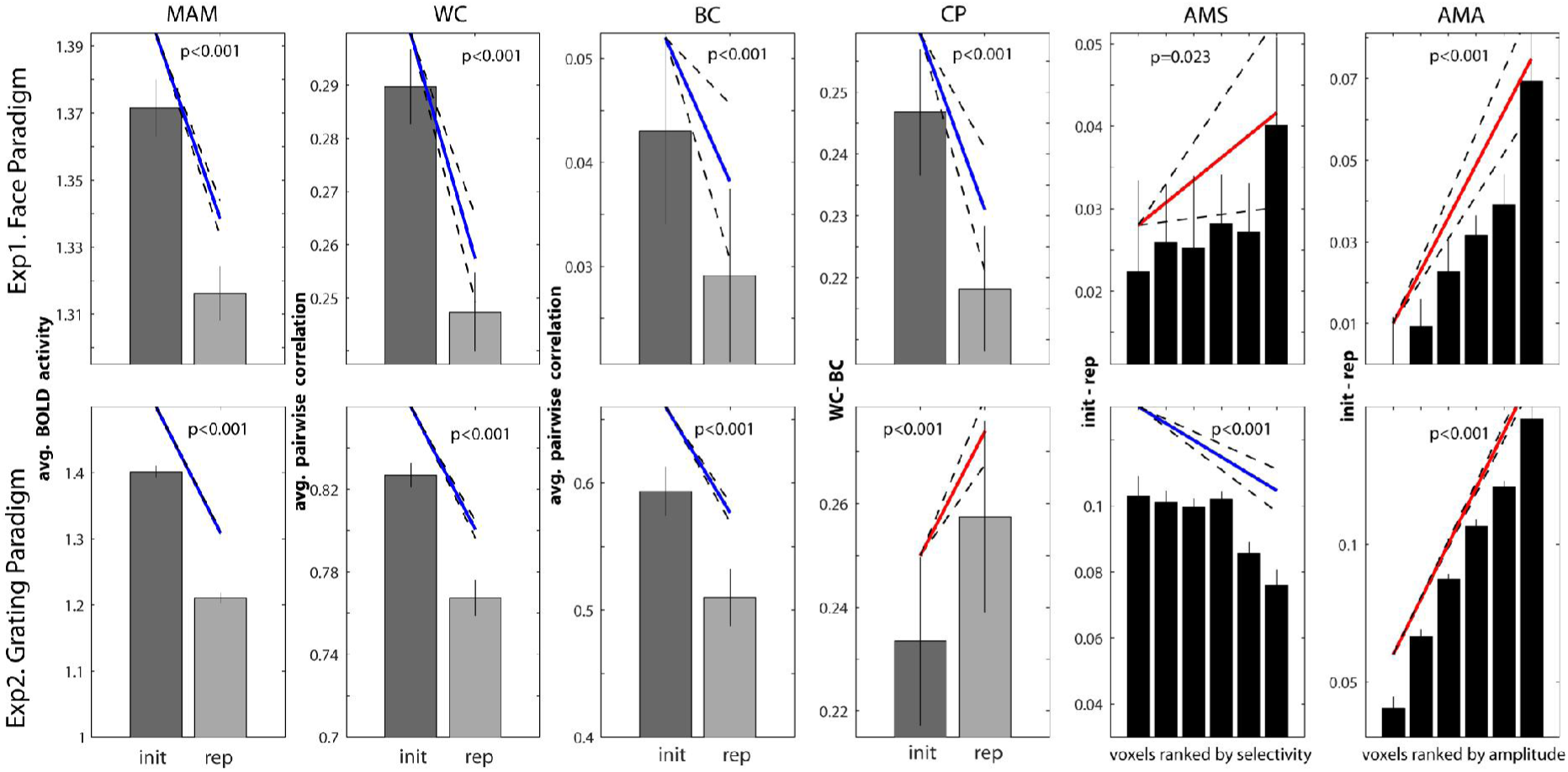
Local scaling prediction for all the 6 criteria after adding correlated neural activity to make BC positive. Around 10% of correlated neural activities (flat curve neurons) were added to the voxels in the face Paradigm simulation, and around 50% added to the grating Paradigm simulation. See the supplementary table1 for all the wining parameter combinations that achieve a qualitatively similar result to the data.

### Supplementary Table

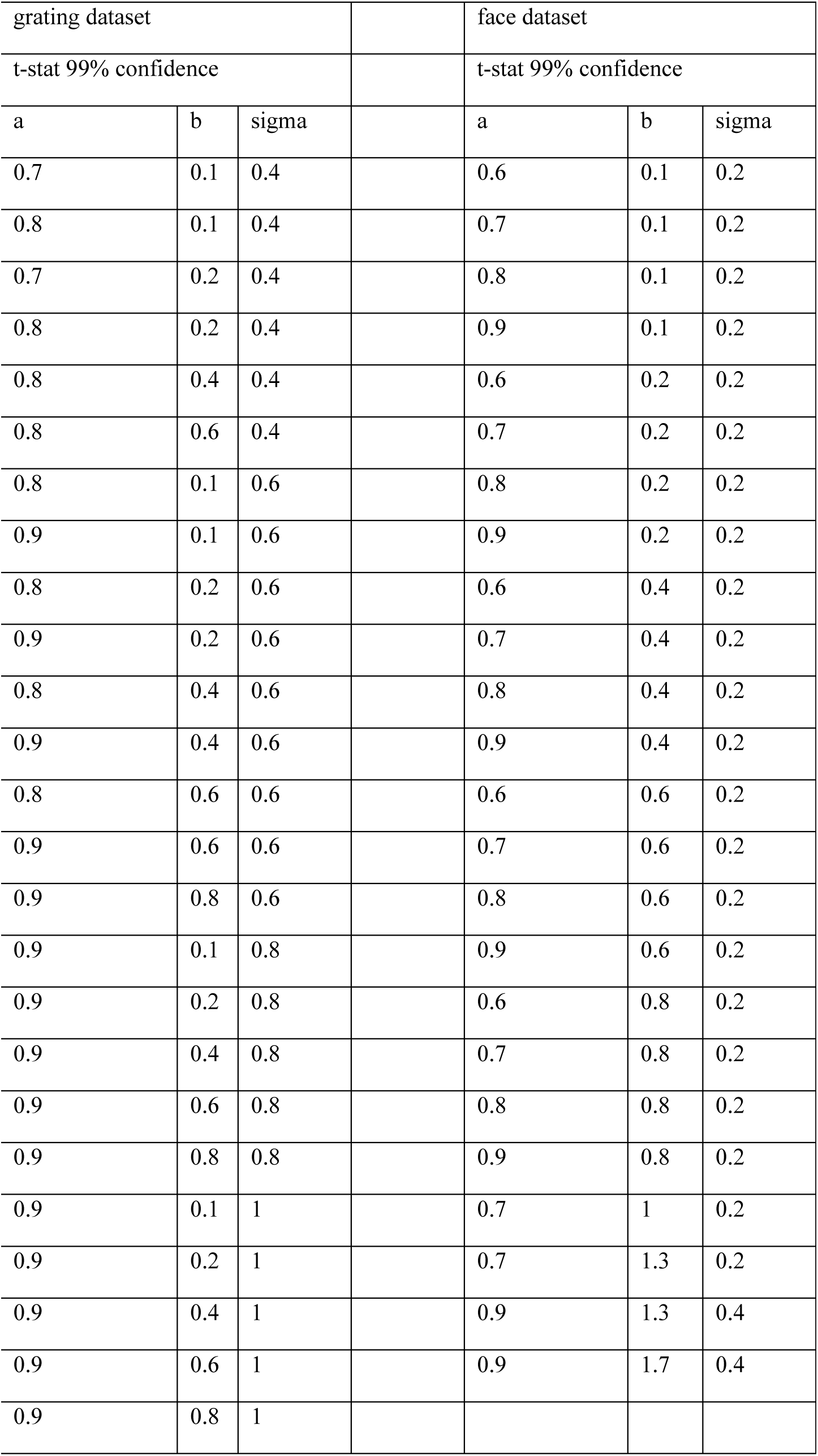

| grating dataset |  |  |  | face dataset |  |  |
| --- | --- | --- | --- | --- | --- | --- |
| t-stat 99% confidence |  |  |  | t-stat 99% confidence |  |  |
| a | b | sigma |  | a | b | sigma |
| 0.7 | 0.1 | 0.4 |  | 0.6 | 0.1 | 0.2 |
| 0.8 | 0.1 | 0.4 |  | 0.7 | 0.1 | 0.2 |
| 0.7 | 0.2 | 0.4 |  | 0.8 | 0.1 | 0.2 |
| 0.8 | 0.2 | 0.4 |  | 0.9 | 0.1 | 0.2 |
| 0.8 | 0.4 | 0.4 |  | 0.6 | 0.2 | 0.2 |
| 0.8 | 0.6 | 0.4 |  | 0.7 | 0.2 | 0.2 |
| 0.8 | 0.1 | 0.6 |  | 0.8 | 0.2 | 0.2 |
| 0.9 | 0.1 | 0.6 |  | 0.9 | 0.2 | 0.2 |
| 0.8 | 0.2 | 0.6 |  | 0.6 | 0.4 | 0.2 |
| 0.9 | 0.2 | 0.6 |  | 0.7 | 0.4 | 0.2 |
| 0.8 | 0.4 | 0.6 |  | 0.8 | 0.4 | 0.2 |
| 0.9 | 0.4 | 0.6 |  | 0.9 | 0.4 | 0.2 |
| 0.8 | 0.6 | 0.6 |  | 0.6 | 0.6 | 0.2 |
| 0.9 | 0.6 | 0.6 |  | 0.7 | 0.6 | 0.2 |
| 0.9 | 0.8 | 0.6 |  | 0.8 | 0.6 | 0.2 |
| 0.9 | 0.1 | 0.8 |  | 0.9 | 0.6 | 0.2 |
| 0.9 | 0.2 | 0.8 |  | 0.6 | 0.8 | 0.2 |
| 0.9 | 0.4 | 0.8 |  | 0.7 | 0.8 | 0.2 |
| 0.9 | 0.6 | 0.8 |  | 0.8 | 0.8 | 0.2 |
| 0.9 | 0.8 | 0.8 |  | 0.9 | 0.8 | 0.2 |
| 0.9 | 0.1 | 1 |  | 0.7 | 1 | 0.2 |
| 0.9 | 0.2 | 1 |  | 0.7 | 1.3 | 0.2 |
| 0.9 | 0.4 | 1 |  | 0.9 | 1.3 | 0.4 |
| 0.9 | 0.6 | 1 |  | 0.9 | 1.7 | 0.4 |
| 0.9 | 0.8 | 1 |  |  |  |  |

